# Fast and robust ancestry prediction using principal component analysis

**DOI:** 10.1101/713172

**Authors:** Daiwei Zhang, Rounak Dey, Seunggeun Lee

## Abstract

Population stratification (PS) is a major confounder in genome-wide association studies (GWAS) and can lead to false positive associations. To adjust for PS, principal component analysis (PCA)-based ancestry prediction has been widely used. Simple projection (SP) based on principal component loading and recently developed data augmentation-decomposition-transformation (ADP), such as LASER and TRACE, are popular methods for predicting PC scores. However, they are either biased or computationally expensive. The predicted PC scores from SP can be biased toward NULL. On the other hand, since ADP requires running PCA separately for each study sample on the augmented data set, its computational cost is high. To address these problems, we develop and propose two alternative approaches, bias-adjusted projection (AP) and online ADP (OADP). Using random matrix theory, AP asymptotically estimates and adjusts for the bias of SP. OADP uses computationally efficient online singular value decomposition, which can greatly reduce the computation cost of ADP. We carried out extensive simulation studies to show that these alternative approaches are unbiased and the computation times can be 10-100 times faster than ADP. We applied our approaches to UK-Biobank data of 488,366 study samples with 2,492 samples from the 1000 Genomes data as the reference. AP and OADP required 7 and 75 CPU hours, respectively, while the projected computation time of ADP is 2,534 CPU hours. Furthermore, when we only used the European reference samples in the 1000 Genomes to infer sub-European ancestry, SP clearly showed bias, unlike the proposed approaches. By using AP and OADP, we can infer ancestry and adjust for PS robustly and efficiently.

## 1 Introduction

Population stratification (PS) is a major confounder for genetic association analysis (Price et al., 2006), and the adjustment of PS requires the estimation of the ancestry structure among study samples. Principle component analysis (PCA) is a multivariate statistical method which finds the direction of the maximum variability (Jolliffe, 2002). By aggregating information across all genetic markers, PCA has been effective for PS adjustment (Reich et al., 2008). To adjust for PS, PCA can be applied to study data to calculate the principle component (PC) scores, which are regarded as variables of ancestry and can be used as covariates to adjust for. An alternative approach is predicting the PC scores of the study samples by using reference genotyped samples with detailed ancestry information. This prediction-based approach allows not only adjustment for PS but also inference of the ancestry memberships of the study samples.

The standard approach of predicting PC scores is to project the study samples onto the maximum variability direction, called PC loadings. In this paper, we call this approach as simple projection (SP). However, when the number of features greatly exceeds the size of the reference samples, which is common for genome-wide association data, the PC scores predicted by SP are known to be systematically biased toward null (Dey and Lee, 2019). This shrinkage bias can cause inaccurate prediction of the ancestry of each study sample and inappropriate adjustment of PS.

One way of addressing this shrinkage bias is presented by Wang et al. (2014, 2015). Their solution is to combine one study sample with all the reference samples and find the PC scores of this augmented data set. The PC scores of the study individuals are then mapped to the reference sample PC space by a Procrustes transformation. We call this method “augmentation, decomposition, and Procrustes transformation” (ADP). This method has been shown to be effective in eliminating the shrinkage bias of study PC scores. However since ADP needs to run PCA separately for each of the augmented dataset, it is computationally expensive, especially with large reference samples. For example, the estimated computation time for predicting the ancestry of UK-Biobank data of 488,366 samples with 2,492 reference samples is 2,534 CPU hours. Since computation time is cubic to the reference sample sizes, the computation time will rapidly increase with larger reference samples.

To address the limitations of SP and ADP, we develop and propose two alternative methods for the ancestry prediction and applied them to UK-Biobank data. The first approach removes the bias in SP by estimating the asymptotic bias factor, which is calculated based on random matrix theory (Dey and Lee, 2019). The second approach improves the computational efficiency of ADP by using an online singular value decomposition (SVD) algorithm (Halko et al., 2011), which obtains the SVD results of the augmented matrix by updating the SVD results of the reference matrix, since the latter only differs slightly from the former and many of the overlapping calculations can be avoided. We call the first approach “bias-adjusted projection” (AP) and the second approach “online augmentation, decomposition, and Procrustes transformation” (OADP). Through the extensive simulation studies, we have shown that AP and OADP have both achieved accuracy close to that of ADP and computational efficiency close to that of SP. The UK-Biobank data application shows that the proposed approaches are 20-100 times faster than ADP while achieving the similar accuracy.

## 2 Method

### 2.1 Model and PCA on the reference data

For PC score prediction, we have reference and study sample data, which can be represented by matrices. Let *<u>X</u>* be a *p × n* matrix of reference genotypes and *<u>Y</u>* be a *p × m* matrix of study sample genotypes, where *p* is the number of genetic markers, *n* is the number of reference samples and *m* is the number of study samples. In our study we only consider genotypes composed of biallelic single nucleotide polymorphisms (SNP), so each entry of *<u>X</u>* and *<u>Y</u>* is a minor allele count of 0, 1, or 2. For PCA, the reference data matrix is commonly standardized by subtracting the marker mean from each marker genotype and then dividing them by the marker standard deviation. The sample matrix *<u>Y</u>* also can be standardized using marker mean and standard deviation calculated from the reference samples. Suppose *X* and *Y* are the standardized reference and study data matrices, respectively. The sample covariance matrix is *S* = *XX*^*T*^ */n*, and then by eigen-decomposition,

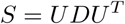

where *D* = diag(*d*_1_, …, *d*_*n*_) is an *n × n* diagonal matrix of ordered sample eigenvalues and *U* = (*u*_1_, …, *u*_*n*_) is a *p × n* corresponding eigenvector matrix. The *j*^th^ PC score vector is 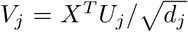, where *U*_*j*_ is the *j*^th^ sample eigenvector, which is also called as the *j*^th^ PC loading. Alternatively PC loading and scores can be calculated using SVD, which is computationally far more efficient when p is much larger than n. By SVD

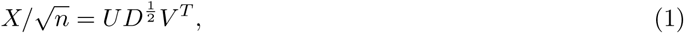

where *V* = (*v*_1_, *…, v*_*n*_) is the left singular vector matrix and *v*_*j*_ is the *j*^th^ PC scores. From (1),

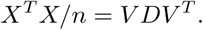

After calculating *v*_*j*_ and *d*_*j*_ from the eigen-decomposition of *X*^*T*^ *X*, the *j*^th^ loading, *u*_*j*_, can be calculated as *U_j_* = *X^T^ V_j_*=*d_j_*.

### 2.2 Predicting the PC scores of the study samples

Here we describe the existing approaches, SP and ADP, and the proposed approaches, ASP and OADP, and their computational complexity to predict *K* top PC scores. Without loss of generality we assume that *K ≪ n ≪ p*.

#### Simple Projection (SP)

SP directly uses the PC loadings of the reference sample PCA to predict the PC scores of the study samples. The algorithm and the computation cost for predicting top *K* PCs are (including steps of reference sample PCA)

1. Perform the reference sample PCA: *X*^*T*^ *X* = *V D*^2^*V* ^*T*^. (CC: *𝒪*[*pn*^2^].)
2. Compute the PC loading matrix for the top *K* PCs: 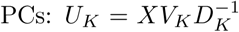. Here *V*_*K*_ and *D*_*K*_ are the the first *K* columns of *V* and the upper-left *K × K* sub-matrix of *D*, respectively. (CC: *𝒪*[*npK*].)
3. Compute the predicted study PC scores for the top *K* PCs: *W*_*K*_ = *Y*^*T*^*U*_*K*_. (CC: *𝒪*[*mpK*].)

The total computational complexity is *𝒪*[*pn*^2^ + *mpK*], which is the lowest among all the methods discussed in this paper. However, though computationally efficient, a major disadvantage of SP is the loss of accuracy when the number of makers, *p*, greatly exceeds the reference sample size, *n*, a situation that is common in GWAS. Lee et al. (2010) have shown that when *n < p*, the predicted PC scores can be shrunken toward the zero. This shrinkage bias limits the accuracy of SP for high dimensional data.

#### Adjusted Projection (AP)

This method calculates the asymptotic shrinkage bias of SP and adjust for predicted PC scores using the estimated bias. The estimation of the bias requires all the eigenvalues of the the reference data matrix. The details for estimating the shrinkage factor are described in Dey and Lee (2019). First, the population eigenvalues are assumed to follow a generalized spiked population model (GSP), where only a few eigenvalues are large (which are called distant spikes) compared to the rest of them. The rest of the eigenvalues are relatively small but not necessarily all equal to each other. Then for the top few PCs that correspond to the distant spikes, the ratio of the population PC scores and the sample PC scores predicted by SP converges in probability to the ratio of the corresponding population eigenvalues (distant spikes) and the sample eigenvalues, as *p → ∞, n → ∞, p/n → γ < ∞*. Dey and Lee (2019) provides two consistent estimators for the distant spikes. One is faster when the number of distant spikes is known, while the other is useful when such information is unknown.

The method for approximating the shrinkage factor has been implemented in the hdpca package in the R language (Dey and Lee, 2016). The algorithm of AP is summarized below.

1. Perform the reference sample PCA and estimate the shrinkage factor *f*_*j*_, (*j* = 1, …, *K*) for each of the top *K* PCs. (CC: *𝒪*[*pn*^2^].)
2. Compute the PC loading matrix for the top *K* PCs with the adjustment for the shrinkage bias: 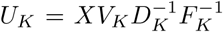. Here *F*_*K*_ = *diag*(*f*_1_, *…, f*_*K*_), is *K × K* diagonal matrix of K shrinkage factors. (CC: *𝒪*[*pnK*].)
3. Compute the predicted study PC scores for the top *K* PCs: *W*_*k*_ = *Y* ^*T*^ *U*_*k*_. (CC: *𝒪*[*mpk*].)

The total computational complexity is *O*[*pn*^2^ + *mpk*], which is the same as that of SP. This is because shrinkage factor estimation is asymptotic-based and can be done rapidly with the sample eigenvalues. In addition, the shrinkage factor only needs to be calculated once for all the study samples.

#### Augmentation, Decomposition, and Procrustes Transformation (ADP)

ADP, such as LASER and TRACE (Wang et al., 2014, 2015), predicts the study PC scores by using a different approach compared to SP and AP. ADP first augments the (standardized) reference matrix by one column by appending a (standardized) study sample. Then SVD is applied to the *p ×* (*n* + 1) augmented matrix 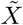 The resulted (*n* + 1) *×* (*n* + 1) normalized PC score matrix 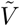 can be divided into two submatrices: the first *n* rows 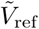, which correspond to the reference samples, and the last row 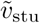, which corresponds to the study sample. Since 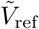 is different from *V*, the *n × n* normalized PC score matrix of the reference data, ADP uses the Procrustes transformation to map 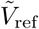 to the original PC space of reference samples. In particular, it consider the following linear transformation,

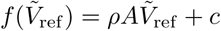

where *ρ* is a non-negative scaling factor, *A* is an *n ×* (*n* + 1) orthonormal matrix for rotation, and *c* is an *n ×* 1 vector for location shift, so that the distance between *V* and the transformed 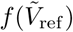 is minimized. We then apply this transformation to 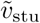 to obtain the predicted PC score, 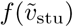. The algorithm is summarized as follows.

1. Perform the reference sample PCA. (CC: *𝒪*[*pn*^2^].)
2. For a study sample *y*, obtain 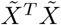 by computing *X*^*T*^ *y*, (*X*^*T*^ *y*)^*T*^, and *y*^*T*^ *y* and appending them to the bottom edge, right edge, and bottom-right corner of *X*^*T*^ *X*. (CC: *𝒪*[*pn*].)
3. Apply eigendecomposition on 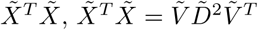. (CC: *𝒪*[*n*^3^].)
4. Find the Procrustes transformation from the first *K′* columns of 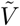 to the first *K* columns of *V*. Note that *K’ > K*. (CC: *𝒪*[*nK*^*t*2^])
5. Go to step 3 for the next study sample until all the study samples are analyzed.

The total computational complexity is *𝒪*[*pn*^2^ + *m*(*np* + *n*^3^)] given that *K*^*t*^ *≪ n ≪ p*. In our study of genotype data, the top four PCs can already seperate the ancestry groups well. We found that for *K* = 4, it is sufficient to let *K′* = 20 for providing accurate prediction of PC scores.

ADP is an nonparameteric approach which does not requires any assumption on the distribution of eigenvalues, and hence can be more robust than AP. It does not suffer the shrinkage bias. The major disadvantage of ADP is its high computational cost. In particular, as the reference size increases, the computational cost for a study sample increases cubicly.

#### Online Augmentation, Decomposition, and Procrustes Transformation (OADP)

Since the augmented data matrix 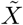 differs in one column only from the reference matrix *X*, the computational process for the SVD of 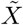 is numerically close to that for the SVD of *X*. If we avoid the repeated computation and obtain the SVD of 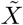 by updating the SVD of *X*, the computation cost can be greatly reduced. One of such “online” algorithms for SVD was proposed for imaging processing (Brand, 2002). This algorithm calculates SVD in the incremental manner and has the ability to rapidly update top few singular values and vectors. Here we propose to use this online SVD to replace the standard SVD for ADP, and call it as “online augmentation, decomposition, and Procrustes transformation” (OADP). The algorithm for this method is as follows:

1. Perform the reference sample PCA. (CC: *𝒪*[*pn*^2^].)
2. Calculate 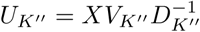 for the first *K″* columns only. (CC: *𝒪*[*K”np*].)
3. 3. Calculate

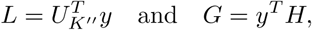

where *H* is the normalized *y − U*_*K*_*″ L*. (CC: *𝒪*[*K″p*].)
4. Calculate *Q*^*T*^ *Q*, where

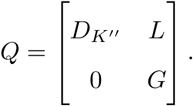

(*𝒪*[*K″*^3^].)
5. Apply eigendecomposition to *Q*^*T*^ *Q* to get 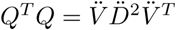. (CC: *𝒪*[*K″*^3^].)
6. Calculate

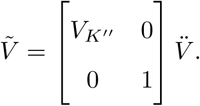

(CC: *𝒪*[*nK″*^2^].)
7. Find the Procrustes transformation from the first *K′* columns of 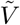 to the first *K* columns of *V*. Note that *K″ > K′ > K*. (CC: *𝒪*[*nK′*^2^])
8. Go to step 3 for the next study individual.

The total computational complexity is *𝒪*[*n*^2^*p* + *m*(*K″p* + *K′*^2^*n*)] provided *K*^*″*^ *≪ n ≪ p*. In our study, we found that setting *K″* = 2*K′* is sufficient for the online SVD method to provide reasonably close approximations to the results of regular SVD algorithms. The computational complexity of OADP for the study individuals increases linearly with respect to the reference sample size, which is much more efficient than ADP’s cubic increasing rate. The closeness between the results given by OADP and ADP will be empirically shown in Section 3.

A comparison of computational complexities of the four PCA methods are shown in Table 1.

**Table 1:** Comparison of computational complexity for different methods. Here *p* is the number of SNPs, *n* is the reference sample size, and *m* is the study sample size. For OADP, the top *K*^*″*^ PCs are computed by the online SVD algorithm, but only the top *K*^*′*^ (*K*^*′*^ *< K*^*″*^) PCs are used by the Procrustes analysis. For SP and AP, the top *K* PCs of the reference samples are computed, and the study samples are projected onto them.

A Tables and Figures
| Method | Reference Complexity | Study Complexity |
| --- | --- | --- |
| ADP | $\mathcal{O}[n^2p]$ | $\mathcal{O}[m(np + n^3)]$ |
| OADP | $\mathcal{O}[n^2p]$ | $\mathcal{O}[m(K''p + K'^2n)]$ |
| SP | $\mathcal{O}[n^2p]$ | $\mathcal{O}[mKp]$ |
| AP | $\mathcal{O}[n^2p]$ | $\mathcal{O}[mKp]$ |

### 2.3 Simulation Studies

We simulated the genotype data using a coalescence-based grid simulation approach by Mathieson and McVean (2012) with population migration. In this approach, we simulated 4 different population groups and hence an 2 *×* 2 grid was created. In each population, we generated (*n* + *m*)*/*2 haploid genotypes with 100,000 biallelic genetic markers. Then we combined every two of the haploid genotypes to form (*n* + *m*)*/*4 duploid genotypes in each population. We used a large migration rate, M=100, to evaluate the performance of the proposed and existing methods in fine-scale population differentiation. Among the (*n* + *m*) generated samples, we randomly selected reference and study samples. The reference sample size *n* ranged from 1000 to 3000, and the study sample size *m* was fixed to 200.

After the individuals were simulated, we applied SP, ADP, AP, and OADP to the data to predict the PC scores for the study samples. We only calculated the top 2 PCs, and for OADP and ADP, we calculated the top 8 PC scores (i.e. *K′* = 8) for the study samples and projected them to the 2-dimensional reference PC score space with a Procrustes transformation. (For OADP, we actually calculated the top 16 PC scores (i.e. *K″* = 16) but used only the top 8 (i.e. *K′* = 8) in order to ensure the accuracy of the SVD.) Finally, we used the nearest neighbors method with the number of neighbors equal to 20 to predict which ethnic group a study sample comes from.

To evaluate the accuracy of each method, we obtained the population means of the reference PC scores and calculated the mean squared difference between the reference population means and the corresponding study population means. We also obtained the mean squared difference of the ADP-predicted PC scores and the predicted PC scores from the other methods, because ADP is currently the most reliable approach.

For the comparison of computational cost, we applied each method 5 times and obtained the mean of the run times. Note that the runtime for the reference sample PCA, reading and writing files, and predicting the populations of the study samples from their predicted PC scores were not included. For SP, AP, and OADP, we used our FRAPOSA software, which implemented the methods using Python. For ADP, we used the TRACE software by Wang et al. (2015). All the programs were run on a single CPU core.

### 2.4 UK Biobank data analysis

We applied the proposed and existing methods to the UK Biobank data (Sudlow et al., 2015; Bycroft et al., 2018), which consists of 488,366 individuals in the United Kingdom, with 1000 Genomes Project data as references (Consortium et al., 2015). The 1000 Genomes data set we used excludes children, siblings, and second-degree relatives and contains 2492 samples from Europe, Africa, East Asia, South Asia, and America (Table 2). The UK Biobank group used 148,000 high-quality genotyped SNPs for the ancestry inference. Among them 145,000 SNPs are observed in 1000 Genomes data, which are used in our analysis.

**Table 2:** Population and sup-population counts in the 1000 Genomes data. Related individuals have been removed.

| Population | # Samples | Sub-Population | Count |
| --- | --- | --- | --- |
| Africans | 657 | ACB (African Caribbeans in Barbados) | 96 |
|  |  | ASW (Americans of African Ancestry in SW USA) | 61 |
|  |  | ESN (Esan in Nigeria) | 99 |
|  |  | GWD (Gambian in Western Divisions in the Gambia) | 113 |
|  |  | LWK (Luhya in Webuye, Kenya) | 97 |
|  |  | MSL (Mende in Sierra Leone) | 84 |
|  |  | YRI (Yoruba in Ibadan, Nigeria) | 107 |
| Native Americans | 347 | CLM (Colombians from Medellin, Colombia) | 94 |
|  |  | MXL (Mexican Ancestry from Los Angeles USA) | 64 |
|  |  | PEL (Peruvians from Lima, Peru) | 85 |
|  |  | PUR (Puerto Ricans from Puerto Rico) | 104 |
| East Asians | 503 | CDX (Chinese Dai in Xishuangbanna, China) | 92 |
|  |  | CHB (Han Chinese in Beijing, China) | 103 |
|  |  | CHS (Southern Han Chinese) | 105 |
|  |  | JPT (Japanese in Tokyo, Japan) | 104 |
|  |  | KHV (Kinh in Ho Chi Minh City, Vietnam) | 99 |
| Europeans | 498 | CEU (Utah Residents with N. & W. European Ancestry) | 95 |
|  |  | FIN (Finnish in Finland) | 99 |
|  |  | GBR (British in England and Scotland) | 90 |
|  |  | IBS (Iberian Population in Spain) | 107 |
|  |  | TSI (Toscani in Italia) | 107 |
| South Asians | 487 | BEB (Bengali from Bangladesh) | 86 |
|  |  | GIH (Gujarati Indian from Houston, Texas) | 102 |
|  |  | ITU (Indian Telugu from the UK) | 102 |
|  |  | PJL (Punjabi from Lahore, Pakistan) | 96 |
|  |  | STU (Sri Lankan Tamil from the UK) | 101 |
| Total | 2492 |  |  |

After predicting PC scores using all the reference samples, we predicted the global ancestry membership using the nearest neighbor method. For each of the study sample, its 20 nearest reference samples, as measured by the Euclidean distance between the PC scores, gave a vote to predict its ancestry, where the weight of each neighbor is inversely proportional to its distance to the study sample. Then, we further investigated the subpopulation ancestry using subpopulation-specific reference samples. For example, to investigate the ancestry within the Europeans, we ran the analysis with the 498 European reference samples only.

Since ADP is very slow, we did not apply ADP to all the samples. Instead, we randomly selected 500 study samples and used them to compare ADP and the other methods. The other three methods, SP, AP, and OADP, were applied to to all the study samples. The same measurements for accuracy and computational cost from the simulation study were applied.

## 3 Results

### 3.1 Simulation studies

We have carried out extensive simulation studies to evaluate the accuracy and computation time of the existing (SP and ADP) and proposed (ADP and OADP) methods. The genotype data with 100,000 genetic markers and 1200 to 3200 individuals were simulated using the grid simulation approach by Mathieson and McVean (2012). We randomly selected 200 individuals as study samples, and the remaining individuals were used as reference samples. In total, five different reference sample sizes (1000, 1500, 2000, 2500, 3000) were considered to investigate the performance of the methods with varying reference sample sizes.

Figure 1 shows PCA plot with 1000 reference samples. It clearly shows that PCA has successfully clustered four different groups. As expected, the PC scores predicted by SP were biased toward zero, but no other methods showed the bias, which indicates that AP, ADP and OADP had successfully resolved the bias. As the reference sample sizes increased (Figures S2 to S5), the bias in SP was reduced, but still visible even when reference sample size was 3000.

**Figure 1:**
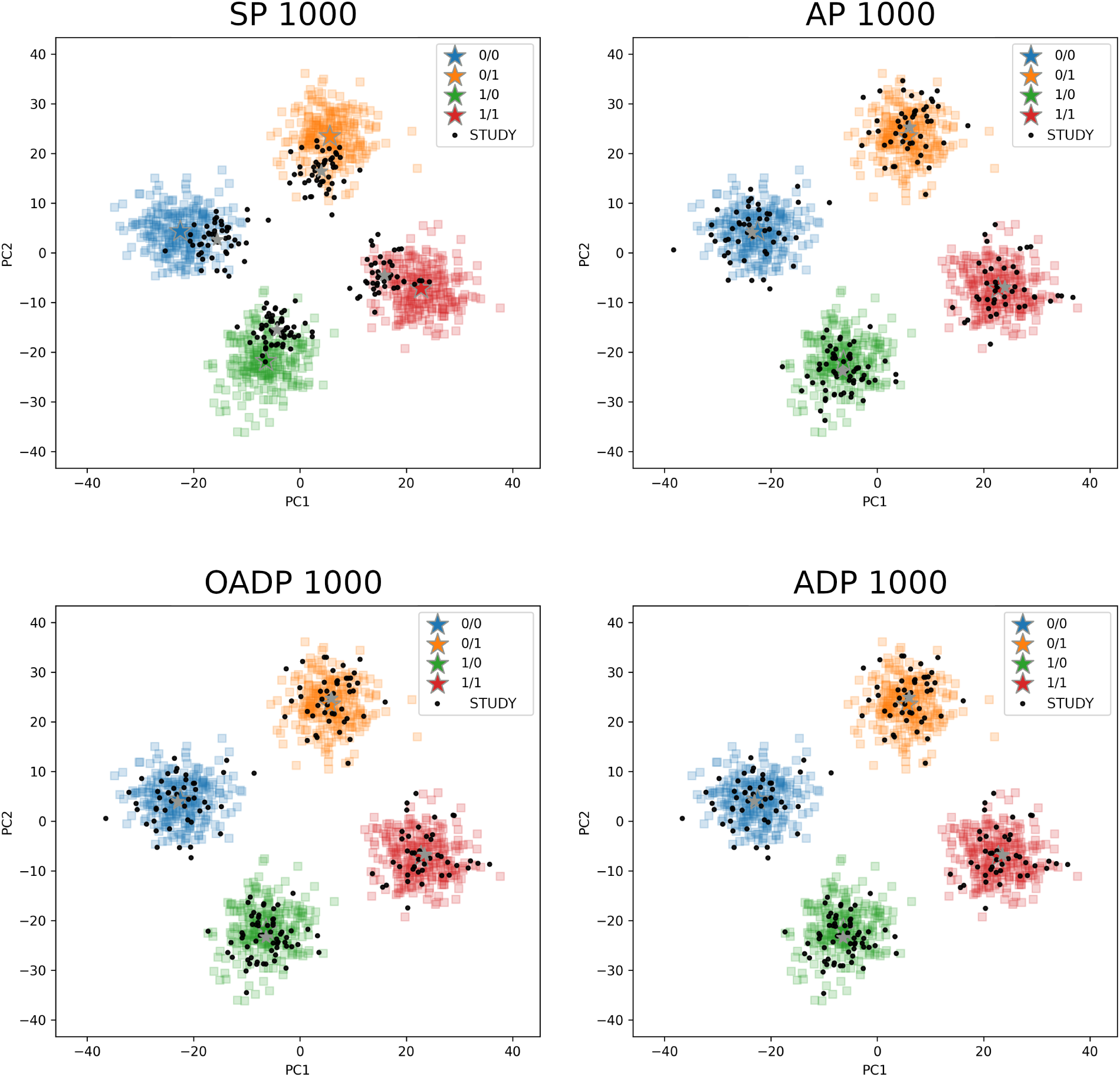
The PC scores predicted by SP, AP, OADP, and ADP when the reference size is 1000. Reference samples are in colors and study samples are in black. The stars mark the mean of each population for the reference and study samples. The number of variants is 100,000, and the study sample size is 200. Only the top 2 PCs are calculated.

To investigate the accuracy of the methods, we obtained the population means of the reference sample PC scores and calculated the mean squared difference (MSD) between the reference population means and the corresponding study population means (Table S1 and Figure 2). As expected, SP had a very high MSD when the reference sample size was 1000 (MSD = 201), and the MSD was reduced as the number of reference samples increased. However, even when the reference sample size was 3000, the MSD was still substantially higher than that of the other methods (MSD = 29 vs. 2.5, 2.2, and 2.4). All the other methods had small MSD regardless of the reference sample sizes. For example, when only 1000 reference samples were used, MSD of AP, OADP, and ADP were only 11, 5, and 6, respectively. Among the proposed approaches, OADP generally had smaller MSD than AP. An interesting observation was that the MSD of ADP were slightly higher than that of OADP (6.2 vs 5.0, 9.7 vs 8.7, 4.2 vs 3.8, 9.6 vs 8.5, and 2.4 vs 2.2 for reference size equal to 1000, 1500, 2000, 2500, and 3000, respectively). We also calculated the MSD between ADP-predicted PC scores and the predicted PC scores from the other methods (ADP-MSD) (Table S1 and Figure S1). As expected, OADP had smaller ADP-MSD than AP.

**Figure 2:**
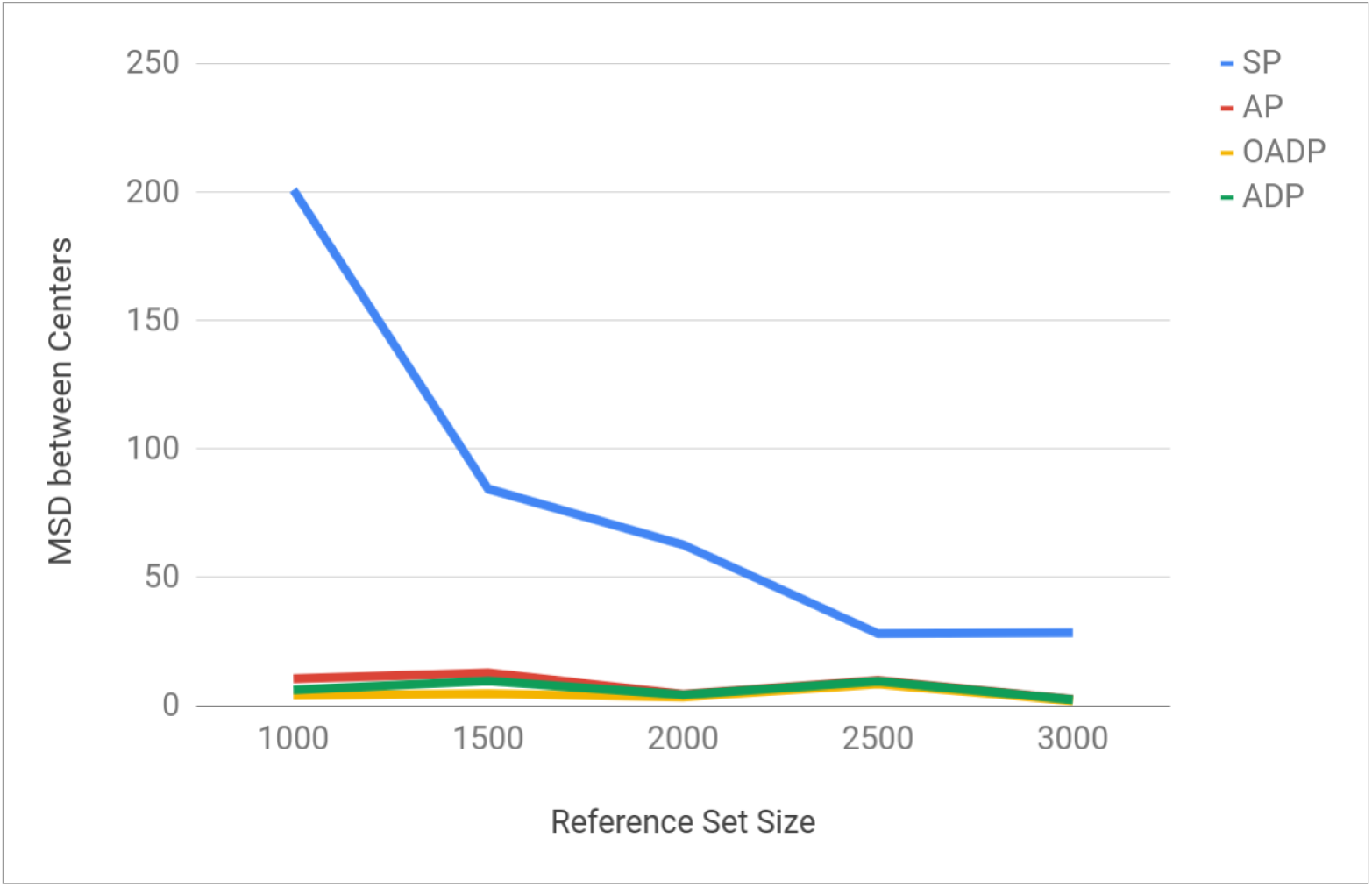
Comparison of the four methods’ accuracy when applied to the simulated data. Accuracy is measured by the mean squared difference between the population means of the reference samples and the corresponding population means of the study samples. The number of variants is 100,000, and the study sample size is 200. Only the top 2 PCs are calculated.

Table S1 and Figure 3 show the computation time of the four methods. ADP had much higher computation time than the other methods, and the computation time of ADP rapidly increased as the reference sample size increased. This is consistent with the theoretical computational complexity of ADP, which is *O*[*n*^3^] (Table 1) when the data dimension and study size are fixed. When the reference sample size reached 3000, ADP’s computation time for predicting 200 study samples was 4487 seconds, which was 38 times higher than that of OADP (119 seconds) and 534 times of SP’s and AP’s (8.4 seconds for both). In a study of 500,000 samples with a reference size of 3000, the projected computation time of ADP would be 3116 CPU hours (130 CPU days). In comparison, the computation time of SP, AP, and OADP increased only slightly with the reference sample size, which is consistent with their *O*[1] and *O*[*n*] computational complexity when the data dimension and study sample size are fixed. For reference sizes between 1000 and 3000, the computation time of SP and AP ranged from 7 to 9 seconds, and that of OADP ranged from 113 to 120 seconds. In a study of 500,000 samples with a reference size of 3000, OADP would only require 82 CPU hours and AP would require 5.8 CPU hours.

**Figure 3:**
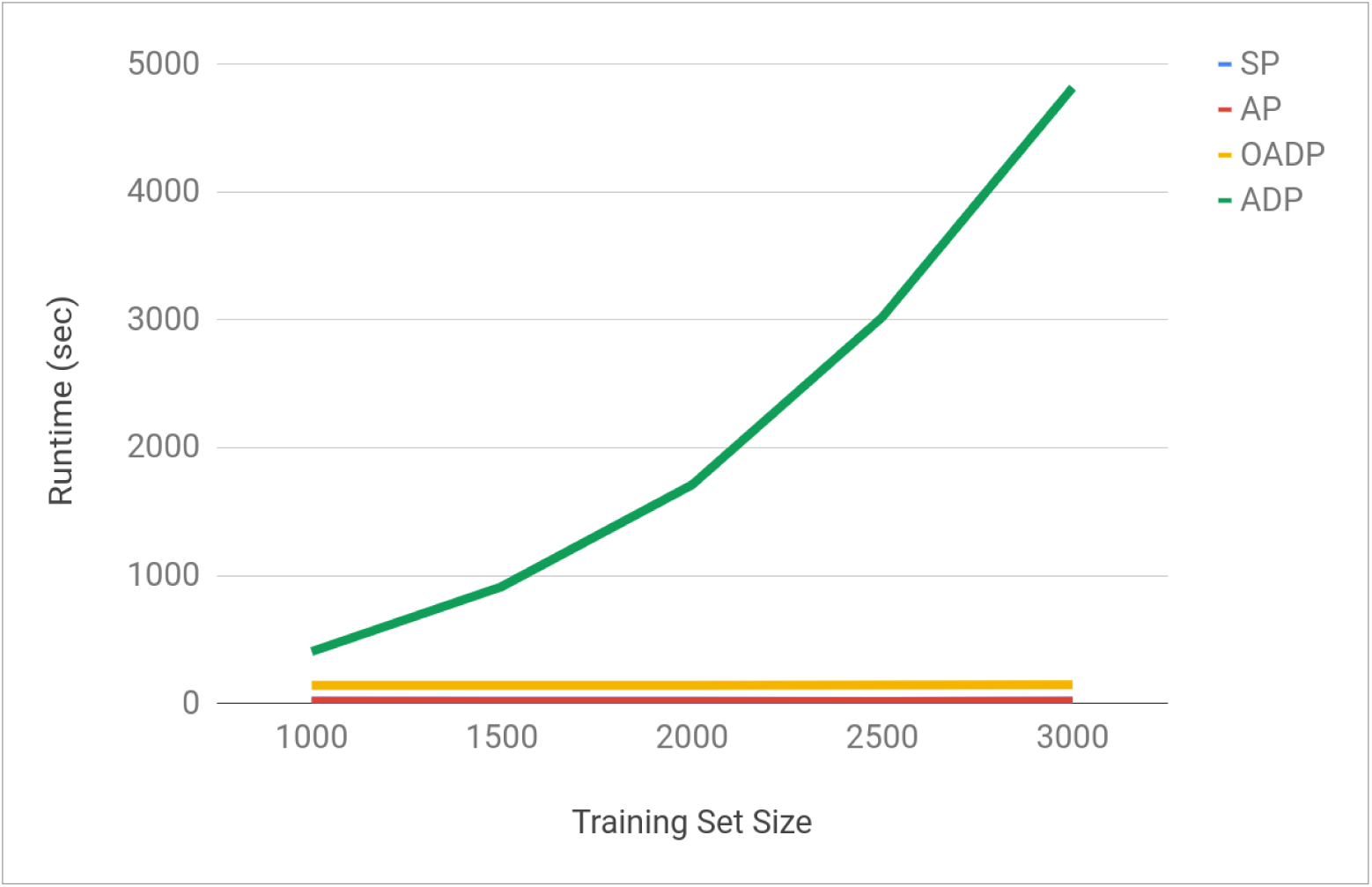
The comparison of the runtimes of the four methods when applied to the simulated data as the reference size increases. The number of variants is 100,000, and the study sample size is 200. Only the top 2 PCs are calculated.

Overall, the simulation study demonstrates that both of the proposed approaches, AP and OADP, are scalable to large sample data and provide accurate PC score prediction. In contrast, the existing approaches suffers from bias or high computation cost.

### 3.2 UK Biobank data analysis

To identify the ancestry structure of the UK biobank data, we applied the proposed and existing approaches by using the 1000 Genomes data as references. The UK Biobank data contains 488,366 samples collected over multiple centers in the United Kingdom. The 2492 independent samples from the 1000 Genomes data were used as the reference set. Sample sizes of the global populations and the sub-ancestry groups are given in Table 2. The predicted populations (by OADP) of the UK Biobank samples are shown in Table 3. Since ADP is computationally too expensive, we only applied ADP to 500 randomly selected samples, which is for the method comparison. All the other methods were applied to all the 488,366 samples.

**Table 3:** Predicted populations of the UK Biobank samples by OADP. Mixed samples are defined to be those whose highest population prediction probability is 0.875 or less by the nearest neighbor method. The number of neighbors is 20, and the weights are inversely proportional to the Euclidean distances.

| Predicted Population | # Samples |
| --- | --- |
| Africans | 8169 |
| Native Americans | 2147 |
| East Asians | 2569 |
| Europeans | 461807 |
| South Asians | 10250 |
| Mixed | 3424 |
| Total | 488366 |

Figure 4 shows the first four PC plots of the reference sample PC scores and the predicted PC scores of the 500 randomly selected UK Biobank samples. It is clear that these four PCs well separated different ancestry groups, and the predicted PC scores from all the methods were nearly identical. There was no shrinkage bias in SP, which may due to the strong differences in the global ancestries. Figure 6 shows SP-, AP- and OADP-predicted PC scores of all the samples. As expected, they produced similar results.

**Figure 4:**
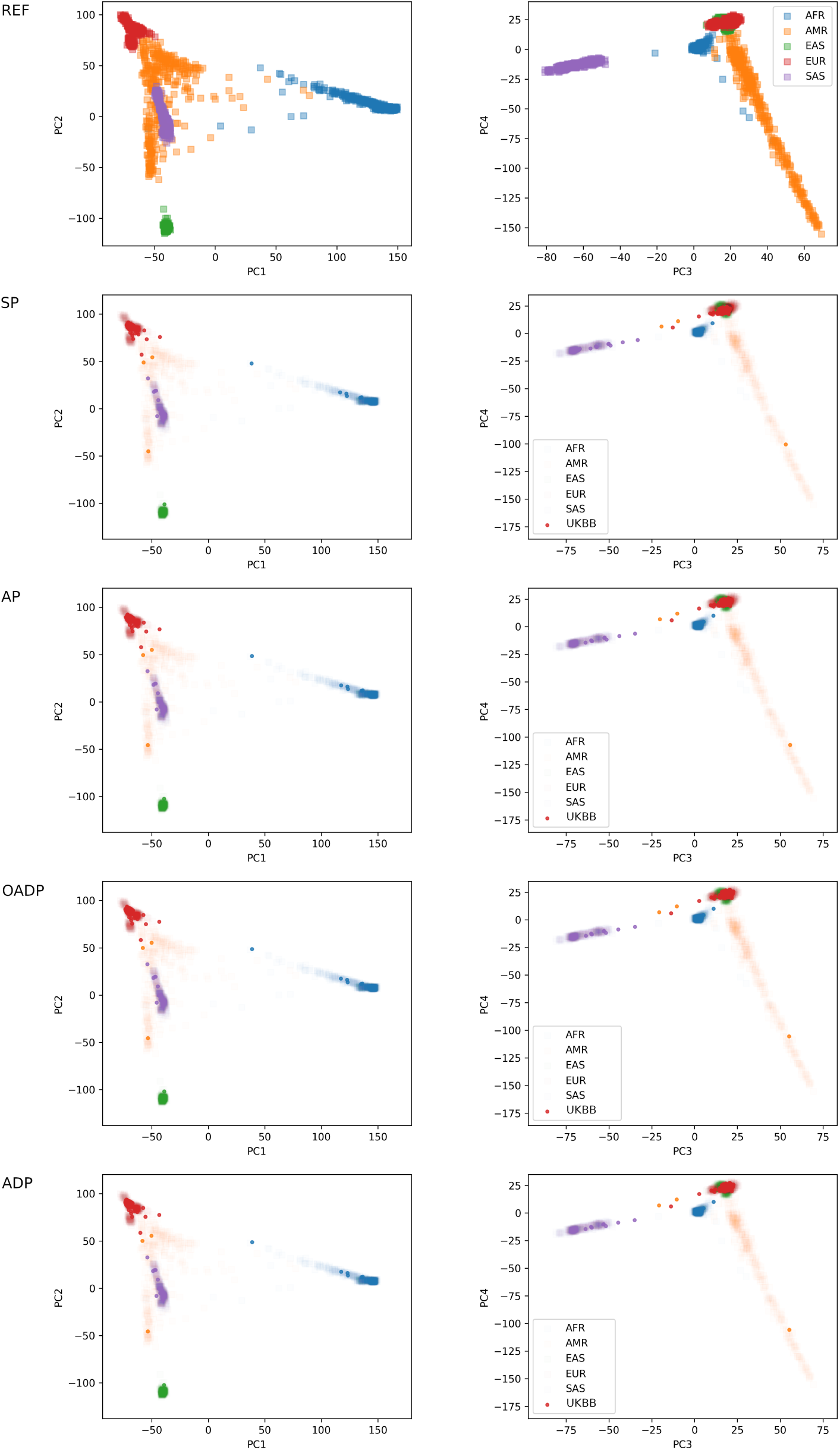
Top row: Reference PC scores of the 2492 global samples from the 1000 Genome data. Bottom four rows: Study PC scores of 500 randomly selected global samples in UK Biobank data predicted by SP, AP, OADP, and ADP. Different colors represent the predicted ancestry of the UK Biobank samples.

**Figure 5:**
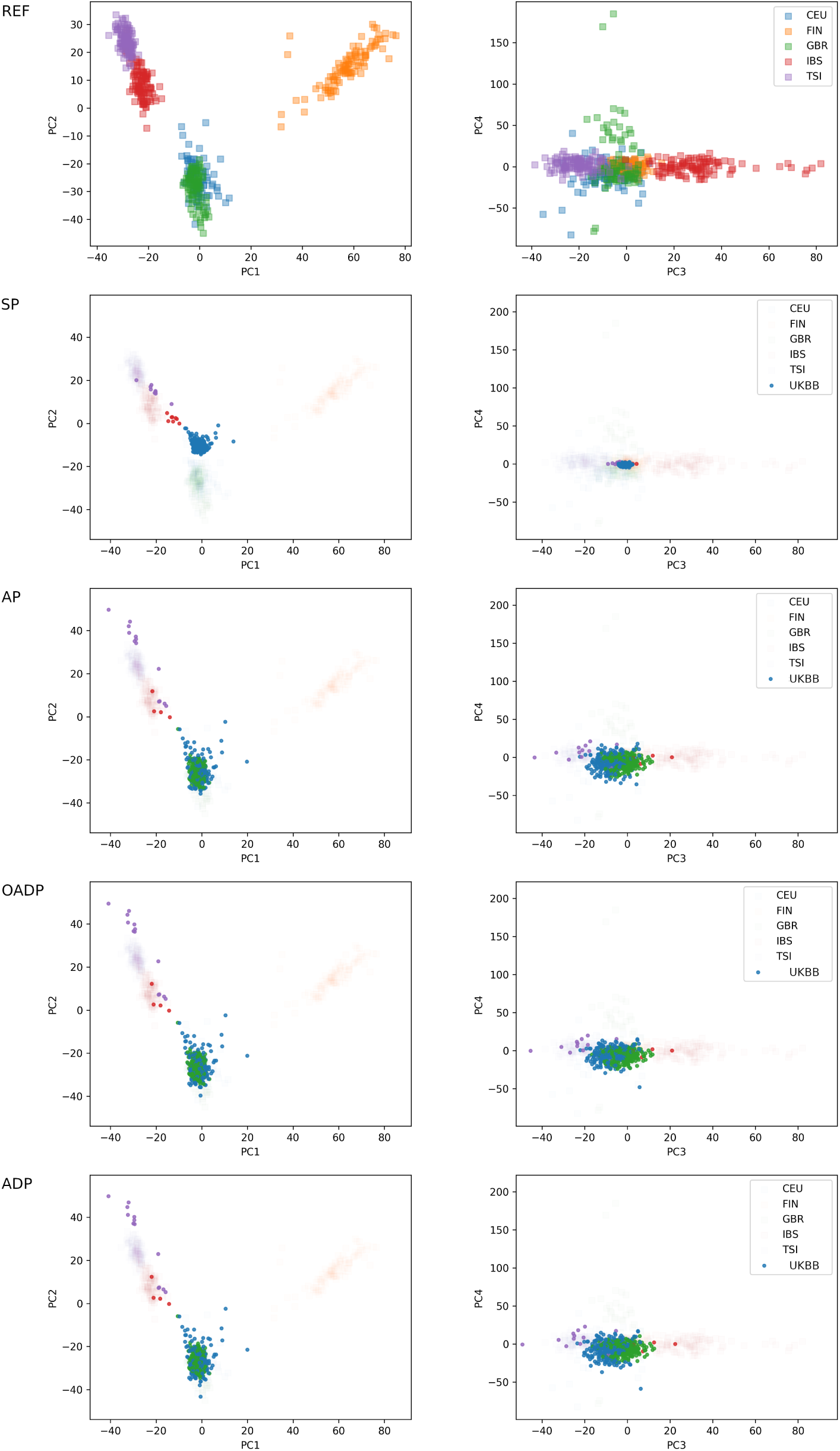
Top row: Reference PC scores of the 498 European samples in the 1000 Genomes data. Bottom four rows: Study PC scores of 500 randomly selected European samples in the UK Biobank data predicted by SP, AP, OADP, and ADP. Europeans samples are those that are predicted to be Europeans by OADP and the nearest neighbor method when using the 2492 global samples from the 1000 Genome data. Different colors represent the predicted ancestry of the UK Biobank samples.

**Figure 6:**
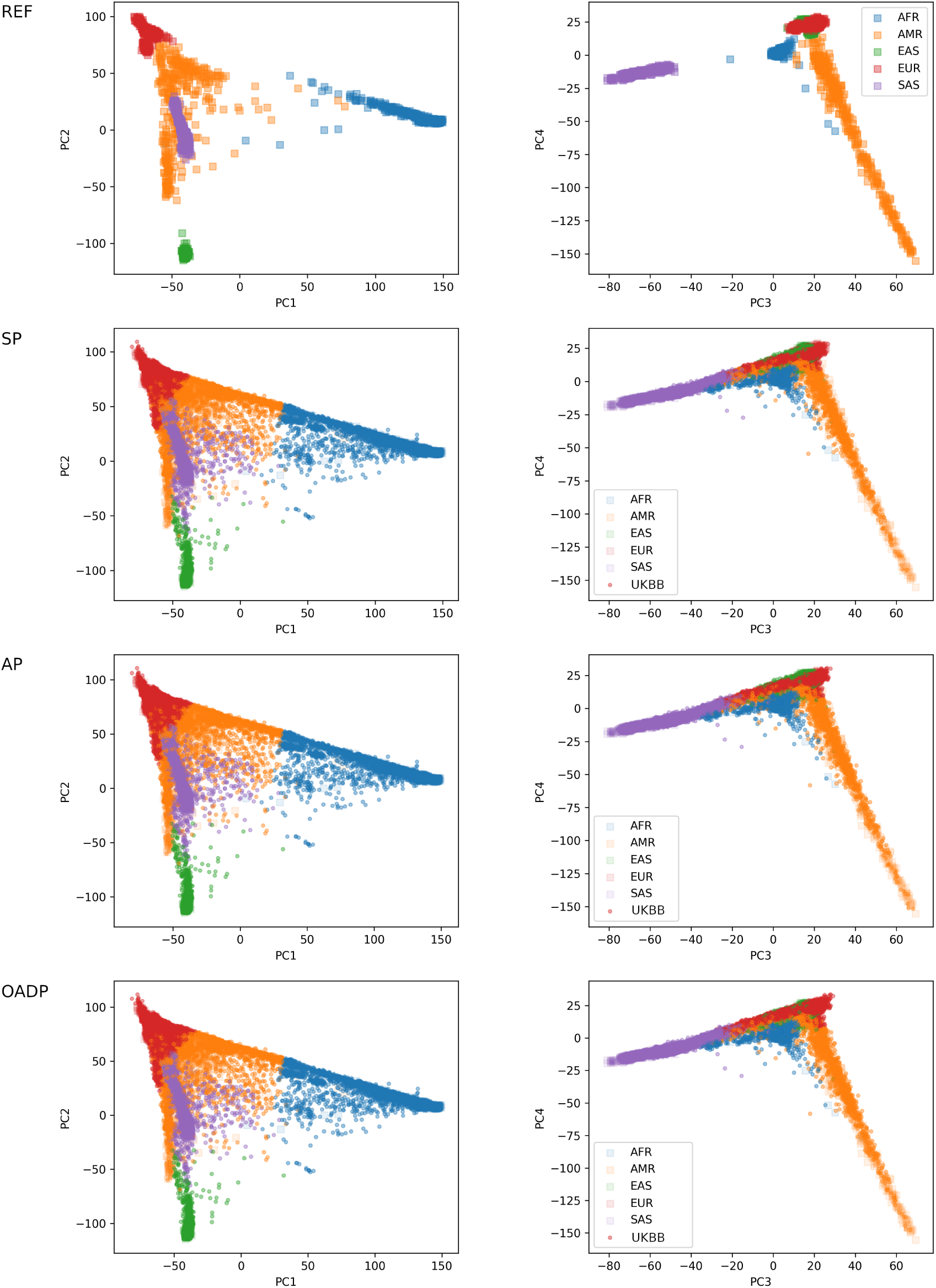
Top row: Reference PC score of the 2492 global samples from the 1000 Genome data. Bottom three rows: PC scores of the 488,366 study samples from the UK Biobank data predicted by SP, AP, and OADP. Different colors represent the predicted ancestry of the UK Biobank samples.

Next, we focused on the sub-ancestries of the European samples. For this, we first identified samples with European ancestry in the UK Biobank using the nearest neighborhood method with the predicted PC scores by OADP. A total of 461,807 samples were identified as European, and we randomly selected 500 of them. Then we predicted the sub-European ancestry of these 500 samples using the 498 European 1000 Genome samples. Figure 5 shows a clear shrinkage bias of SP. Most of the predicted PC scores were shrunken toward zero and nearly none of them were close to the GBR (samples from British in England and Scotland) reference samples. We note that due to similar genetic backgrounds, CEU and GBR reference samples were clustered together. In contrast, AP, ADP and OADP did not show the shrinkage bias, and their predicted PC scores were similar. SP, AP and OADP predicted PC scores of all European ancestry samples were shown in Figure 7. As expected, most of the samples were classified as CEU/GBR by the nearest neighbor method.

**Figure 7:**
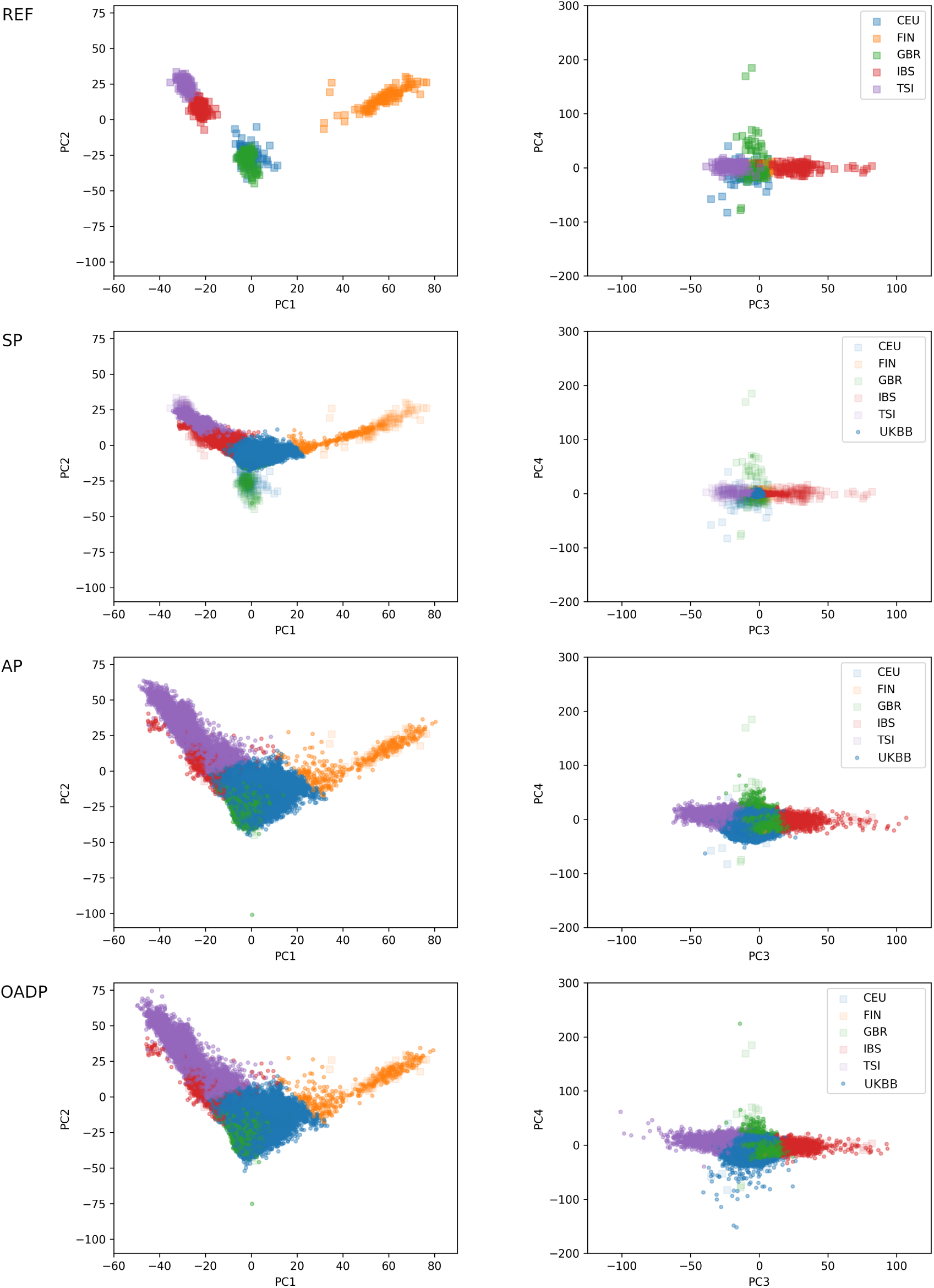
Top row: Reference PC score of the 498 European samples from the 1000 Genome data. Bottom three plots: PC scores of the 461,807 study samples from the UK Biobank data predicted by SP, AP, and OADP. Different colors represent the predicted ancestry of the UK Biobank samples.

We also identified African, East Asian, Native American, South Asian, and mixed-ancestry samples using OADP-predicted PC scores and investigated the sub-ancestry. Mixed samples are defined to be those whose highest population prediction probability is 0.875 or less by the nearest neighbor method. Figures S6 to S9 show the OADP predicted PC scores. One East Asian outlier and three South Asian outliers were removed to make the plots more readable. We note that the PC scores predicted by AP produced similar results (data not shown).

For the computational cost, ADP (401 seconds) needed 1.5 and 15 times more computation time than OADP (275 second) and AP (27 seconds), respectively, for 500 European study samples with 498 European reference samples (Table S2). This difference increased to 33 times (9341 vs 281 seconds) and 346 times (9341 vs 27 seconds) when all the 2492 1000 Genomes reference samples were used as reference. The computation time of SP was almost identical to that of AP. As we included all the 488,366 study samples in the UK Biobank data, AP and OADP only took 7.3 and 77.2 CPU hours, respectively, while ADP was estimated to take 2534 CPU hours.

## 4 Discussion

In this paper, we have compared two existing (SP and ADP) and two novel methods (AP and OADP) of predicting PC scores for the purpose of predicting population structure. The theoretical computational complexity calculation shows that our methods greatly exceed the speed of the existing methods when the reference sample size is large. Moreover, AP improves the the accuracy of SP by adjusting for the shrinkage bias, which is asymptotically estimated by using results from random matrix theory. Our simulation study and the analysis of the UK Biobank data empirically demonstrate the efficiency and unbiasedness of our methods. AP and OADP have been shown to be 10-200 times faster than ADP. They have also successfully separated the sub-populations in the UK-Biobank data when SP shrinks most of the study samples toward zero and is unable to cluster them.

We have noticed a limitation of AP. While the computational complexity and memory usage of SP can be further reduced by using some truncated SVD algorithm (such as the randomized SVD algorithm by Halko et al. (2011)) to compute the SVD for only the top *k* PCs of the reference matrix, AP requires all the eigenvalues and thus a full SVD or eigendecomposition of the reference matrix. This becomes especially important when the reference set is too large. In contrast, OADP needs only the top few singular values and vectors, which can be computed by randomized approaches even for large reference sets.

As the cost of genotyping continues to decrease, the larger genotype data sets will become available. Data with large number of dimension and samples will make it easier to identify and adjust for fine-scale population structure in GWAS, but it also creates a demand for efficient algorithms. When the size of the reference sample set increases, existing methods such as ADP would become impractical to use. But our methods will continue to provide a reasonable computation time and maintain the accuracy and can be a useful tool for genetic studies. The methods have been implemented in the open source software FRAPOSA (github.com/daviddaiweizhang/fraposa).

## Acknowledgement

This work was supported by NIH grant R01 HG008773 (D.Z. and S.L.). It has been conducted using the UK Biobank Resource under application number 45227.

