## Supplementary Materials for "Fast and robust ancestry prediction using principal component analysis"

| Reference Size | 1000 | 1500 | 2000 | 2500 | 3000 |
|---|---|---|---|---|---|
| Runtime (sec) | | | | | |
| SP | 8.1 | 7.7 | 7.9 | 8.8 | 8.4 |
| AP | 8.1 | 8.0 | 8.0 | 8.7 | 8.4 |
| OADP | 113.3 | 114.6 | 114.3 | 119.2 | 118.6 |
| ADP | 410.0 | 803.0 | 1569.2 | 2802.8 | 4487.4 |
| MSD between population means | | | | | |
| SP | 200.9 | 84.4 | 62.7 | 28.1 | 28.5 |
| AP | 10.5 | 12.8 | 4.5 | 9.9 | 2.5 |
| OADP | 5.0 | 8.7 | 3.8 | 8.5 | 2.2 |
| ADP | 6.2 | 9.7 | 4.2 | 9.6 | 2.4 |
| MSD from ADP | | | | | |
| SP | 119.5 | 85.7 | 66.6 | 56.0 | 46.3 |
| AP | 11.5 | 8.3 | 5.0 | 4.7 | 4.2 |
| OADP | 2.4 | 1.5 | 1.1 | 1.7 | 0.5 |
| ADP | 0 | 0 | 0 | 0 | 0 |

Table S1: The study runtimes and errors of the four methods as the reference size increases in the simulation study. The runtimes are the averages of running each setting for five times. "MSD between population means" is the mean squared difference between the means of the reference populations and the means of the study populations. "MSD from ADP" is the mean squared difference between the PC scores predicted by each method and the PC scores predicted by ADP. The number of variants is 100,000, and the study sample size is 200. Only the top 2 PCs are calculated.

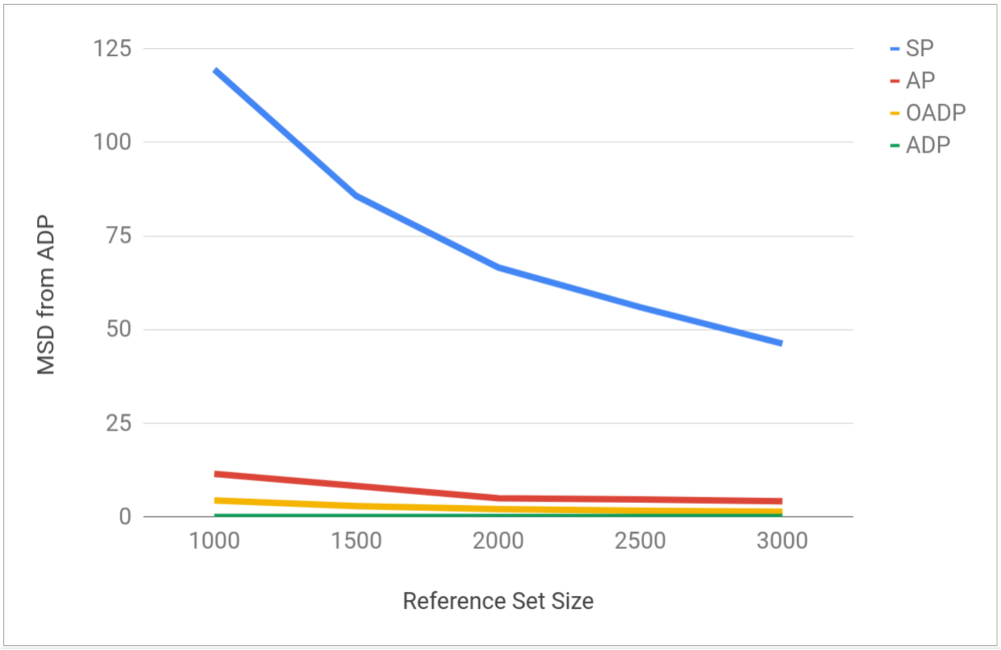


Figure S1: Comparison of the four methods' accuracy when applied to the simulated data, as measured by the mean squared difference between the PC scores predicted by each methods and the PC scores predicted by ADP. The number of variants is 100,000, and the study sample size is 200. Only the top 2 PCs are calculated.

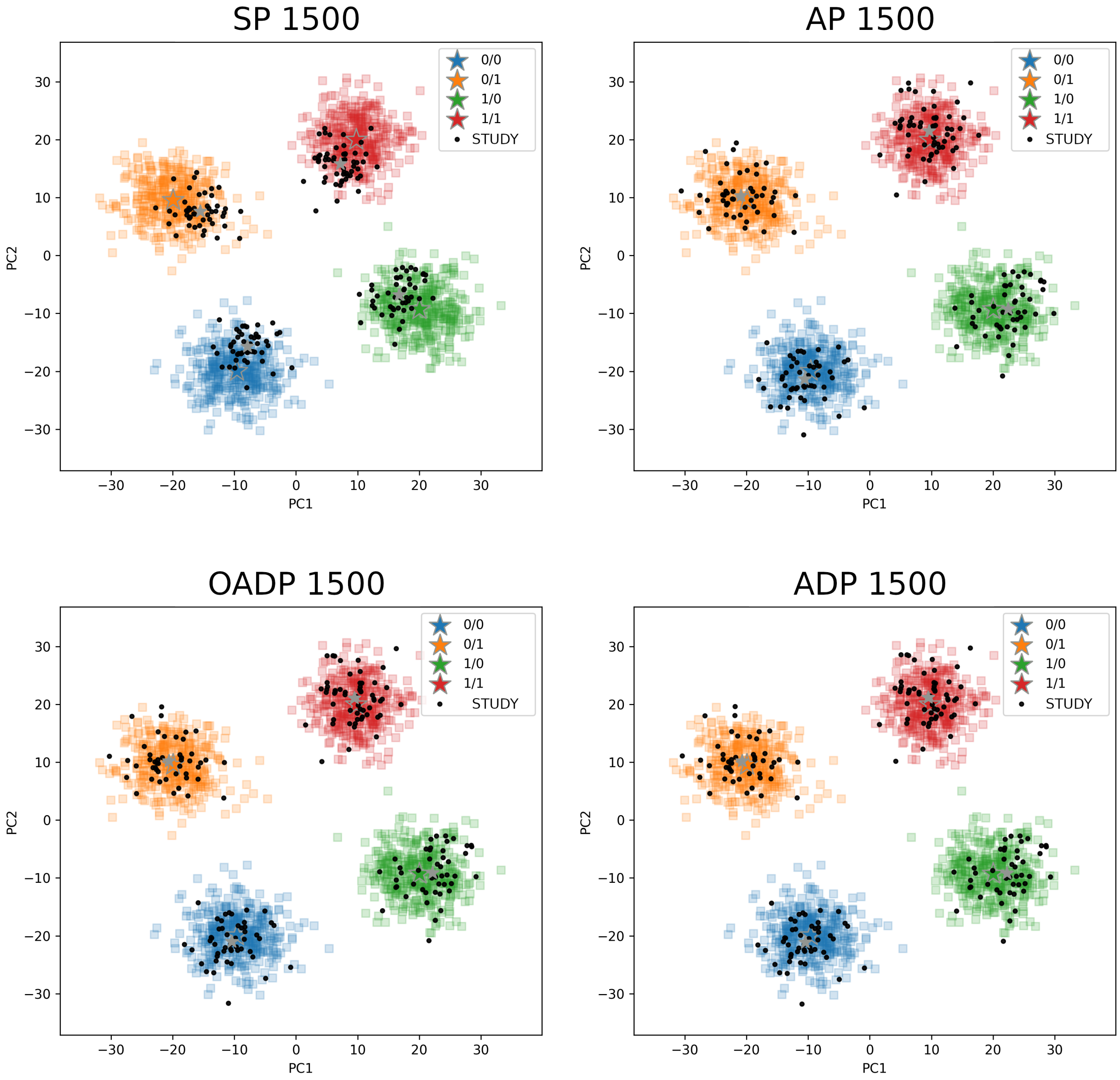


Figure S2: The PC scores predicted by SP, AP, OADP, and ADP when the reference size is 1500. Reference samples are in colors and study samples are in black. The stars mark the mean of each population for the reference and study samples. The number of variants is 100,000, and the study sample size is 200. Only the top 2 PCs are calculated.

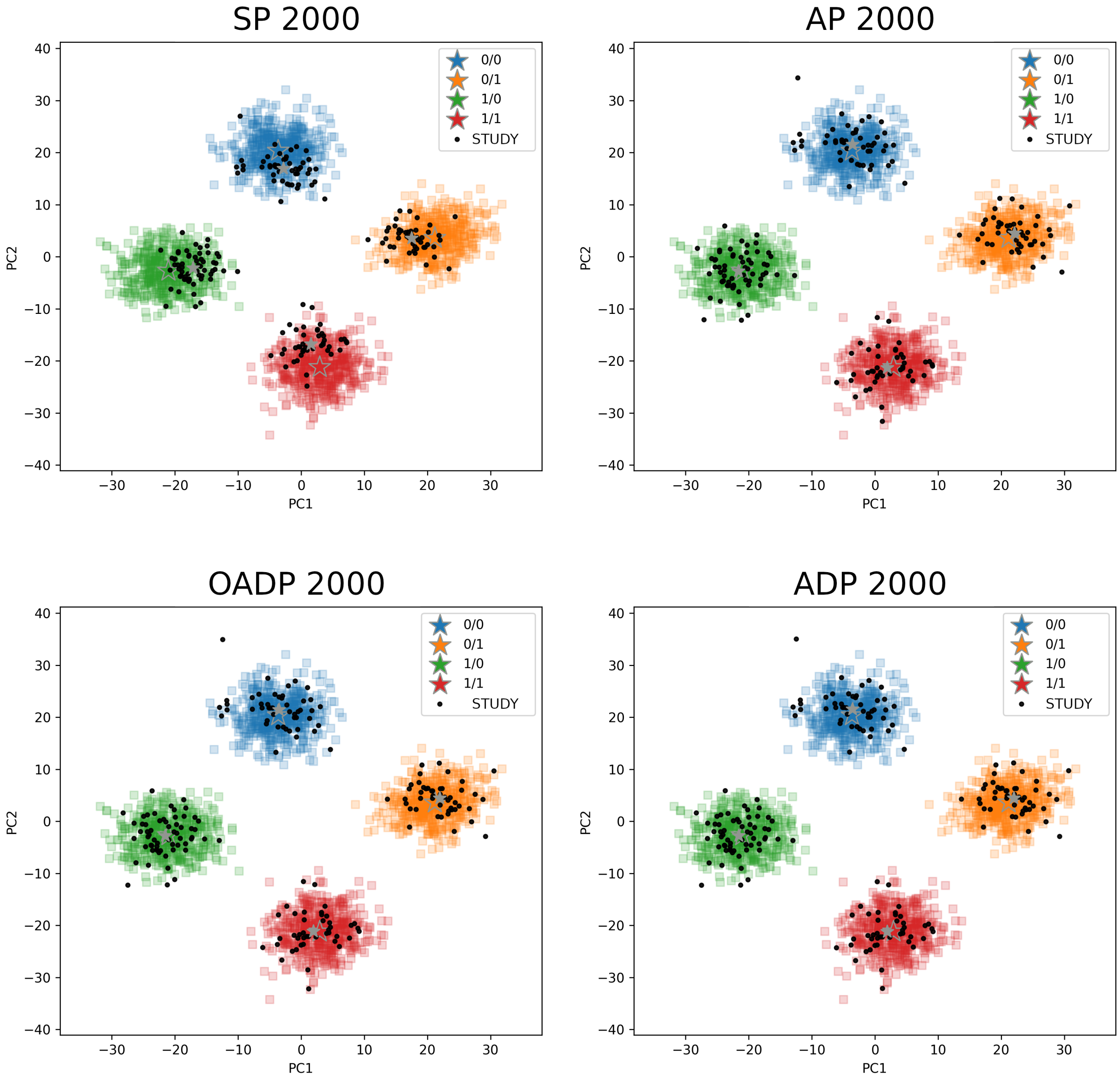


Figure S3: The PC scores predicted by SP, AP, OADP, and ADP when the reference size is 2000. Reference samples are in colors and study samples are in black. The stars mark the mean of each population for the reference and study samples. The number of variants is 100,000, and the study sample size is 200. Only the top 2 PCs are calculated.

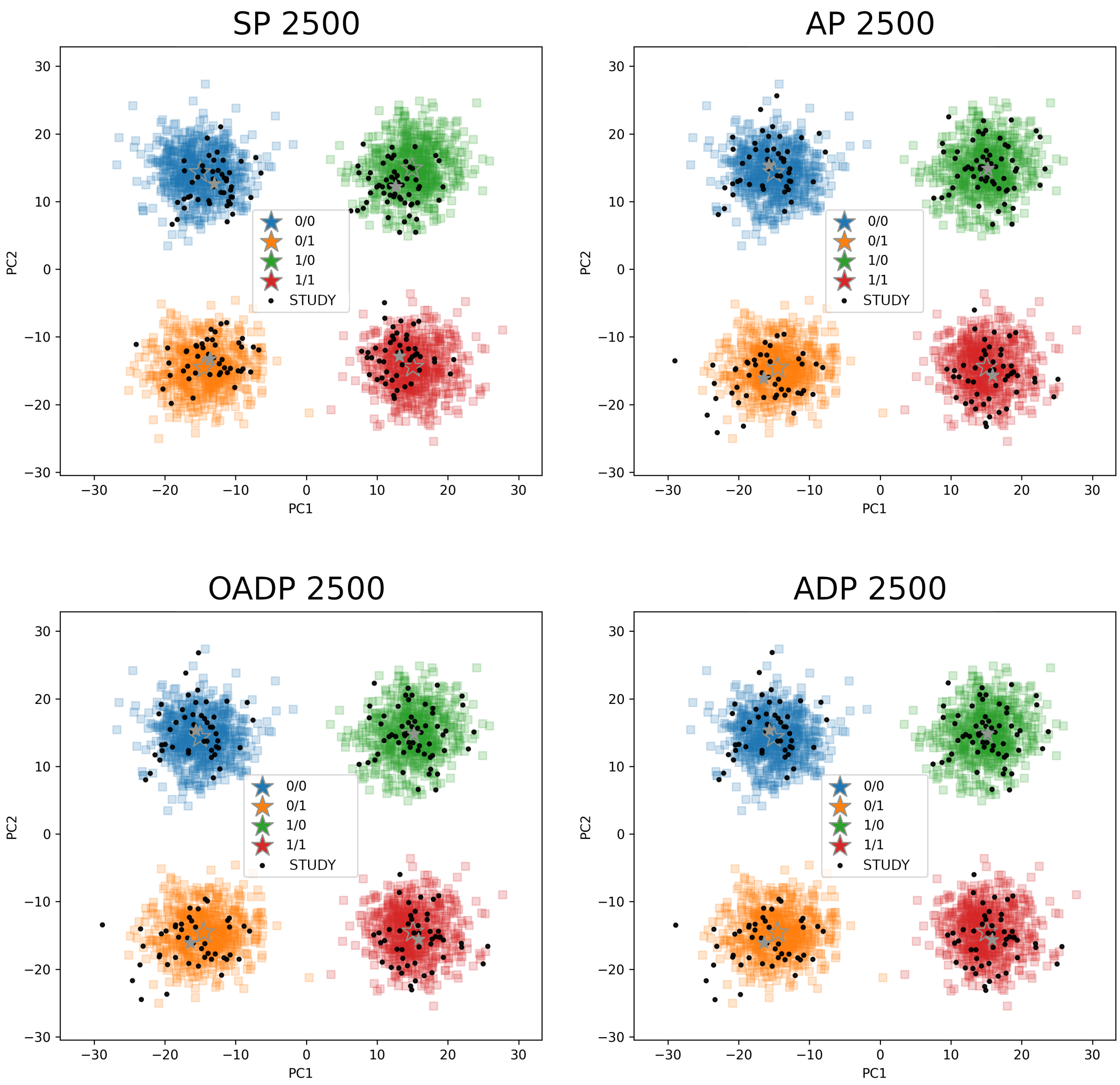


Figure S4: The PC scores predicted by SP, AP, OADP, and ADP when the reference size is 2500. Reference samples are in colors and study samples are in black. The stars mark the mean of each population for the reference and study samples. The number of variants is 100,000, and the study sample size is 200. Only the top 2 PCs are calculated.

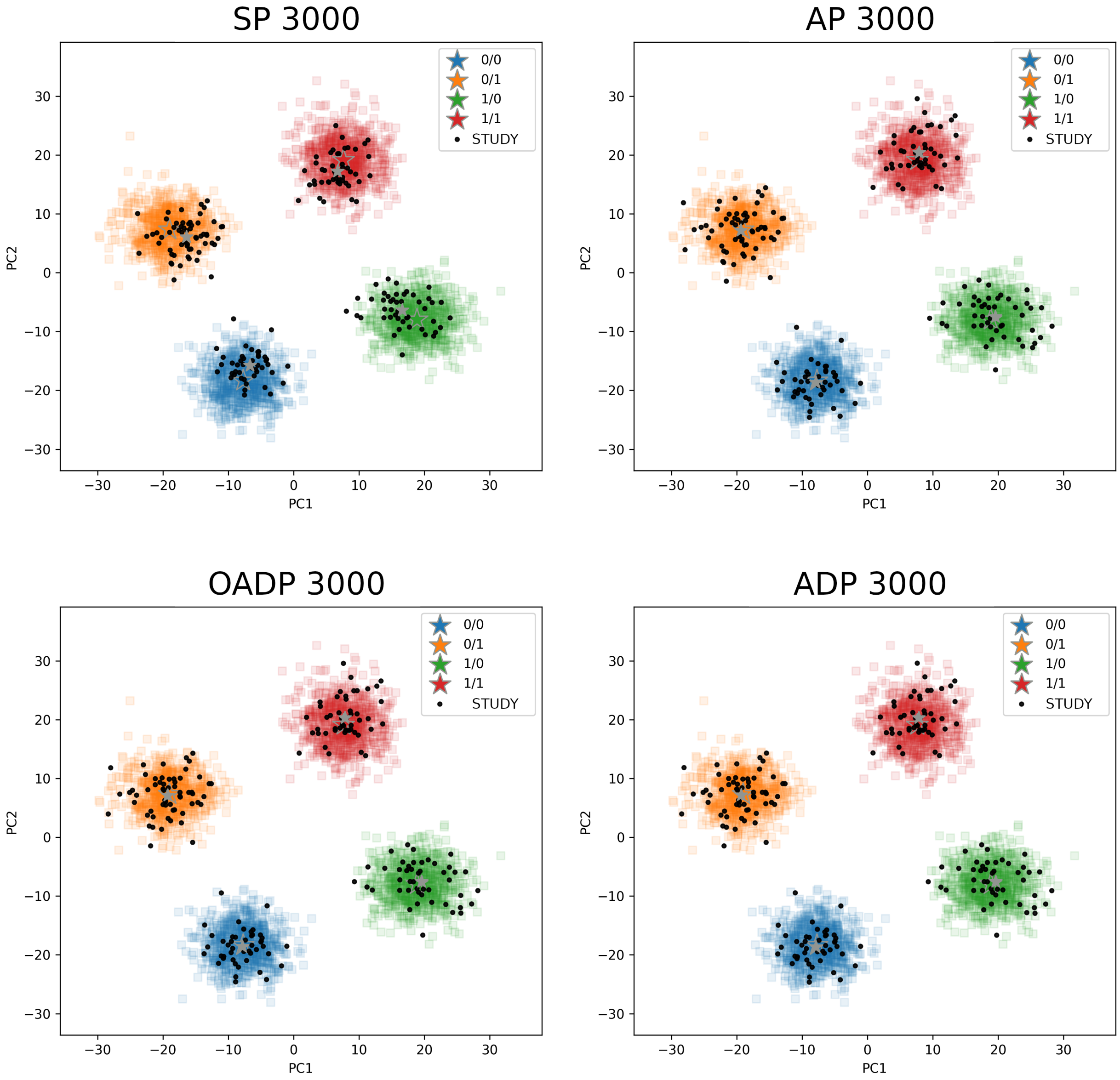


Figure S5: The PC scores predicted by SP, AP, OADP, and ADP when the reference size is 3000. Reference samples are in colors and study samples are in black. The stars mark the mean of each population for the reference and study samples. The number of variants is 100,000, and the study sample size is 200. Only the top 2 PCs are calculated.

| | Runtime (hr) | | Runtime (sec) | | MSD (ADP) | | MSD (popu. means) | |
|---|---|---|---|---|---|---|---|---|
| Population | Global | EUR | Global | EUR | Global | EUR | Global | EUR |
| Ref. size | 2492 | 498 | 2492 | 498 | 2492 | 498 | 2492 | 498 |
| Study size | 488,366 | 461,807 | 500 | 500 | 500 | 500 | 500 | 500 |
| SP | 7.25 | 6.93 | 26.6 | 26.8 | 92 | 473 | 1811 | 1755 |
| AP | 7.28 | 6.95 | 26.7 | 26.9 | 48 | 49 | 1645 | 657 |
| OADP | 77.15 | 72.19 | 281.3 | 274.8 | 10 | 43 | 1719 | 646 |
| ADP | 2534.29* | 102.88* | 9340.8 | 401.0 | 0 | 0 | 1714 | 643 |

Table S2: The study runtimes and MSD for the four methods for 1) all the 488,366 global samples, 2) 461,807 OADP-predicted European samples, 3) 500 randomly selected global samples, and 4) 500 randomly selected OADP-predicted European samples in the UK Biobank data. "MSD (popu. means)" is the mean squared difference between the means of the reference populations and the corresponding means of the study populations. Study populations are predicted by the nearest neighbor methods. "MSD (ADP)" is the mean squared difference between the PC scores predicted by each method and the PC scores predicted by ADP. For the global study samples, the 2492 global samples from the 1000 Genomes data are used as the reference. For the OADP-predicted European study samples, only the 498 European samples from the 1000 Genomes data are used as the reference. The number of variants in common between the reference and study samples is 145,282. Only the top 4 PCs are calculated. *Projected from the runtime of ADP for the 500 global and European samples.

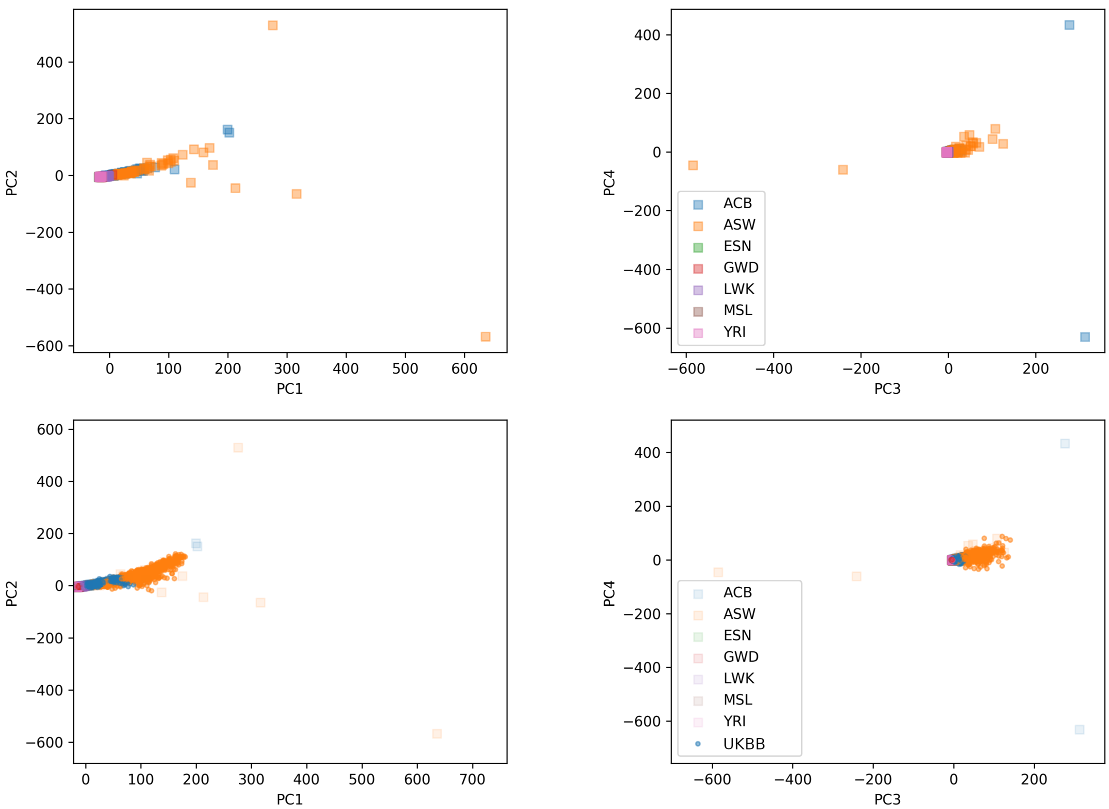


Figure S6: Top plots: Reference PC score of the 657 African samples from the 1000 Genome data. Bottom plots: PC scores of the 8169 African study samples from the UK Biobank data predicted by OADP. Different colors represents the predicted ancestry of the UK Biobank samples.

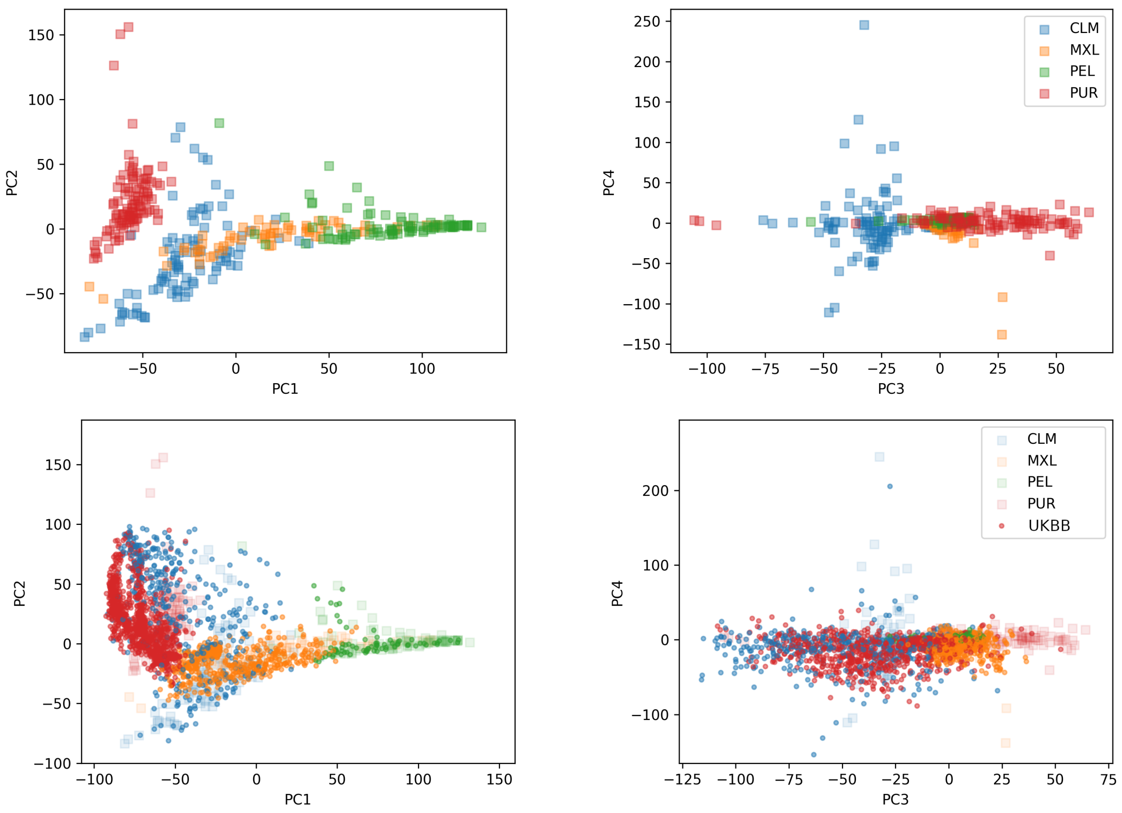


Figure S7: Top plots: Reference PC score of the 347 Native American samples from the 1000 Genome data. Bottom plots: PC scores of the 2147 Native American study samples from the UK Biobank data predicted by OADP. Different colors represents the predicted ancestry of the UK Biobank samples.

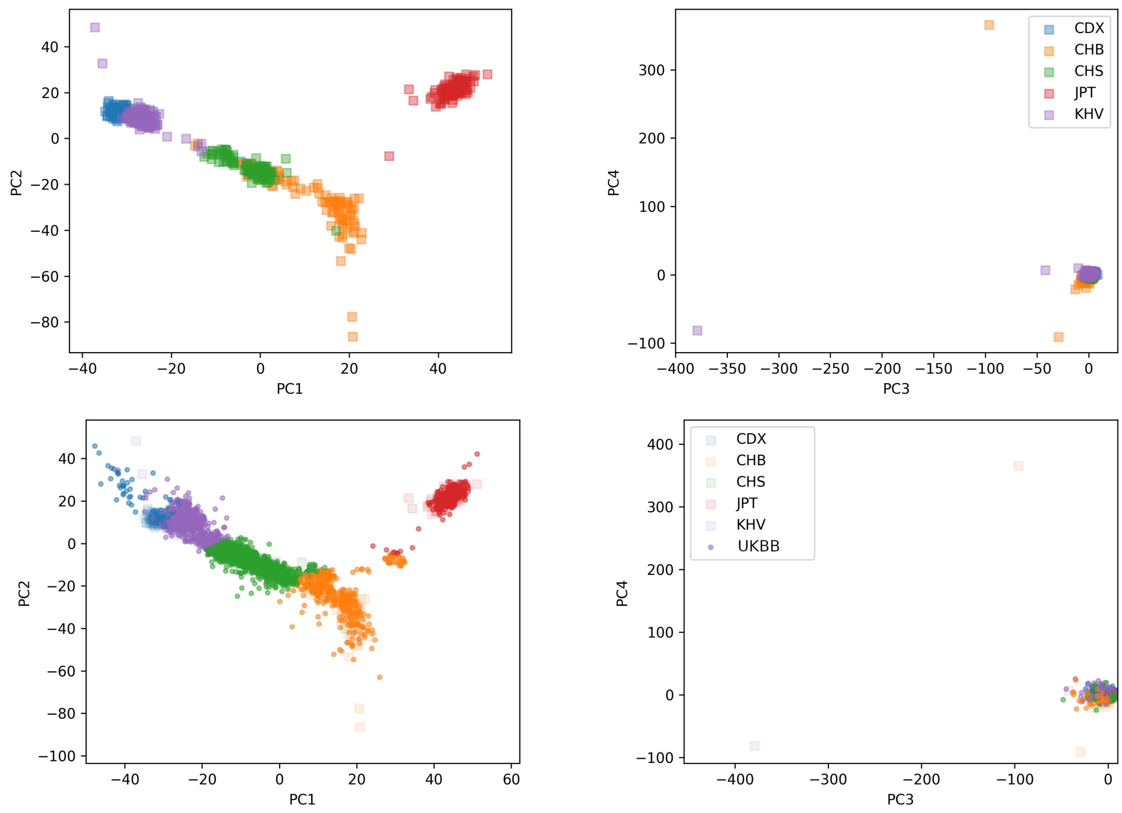


Figure S8: Top plots: Reference PC score of the 503 East Asian samples from the 1000 Genome data. One outlier has been removed. Bottom plots: PC scores of the 2569 East Asian study samples from the UK Biobank data predicted by OADP. Different colors represents the predicted ancestry of the UK Biobank samples.

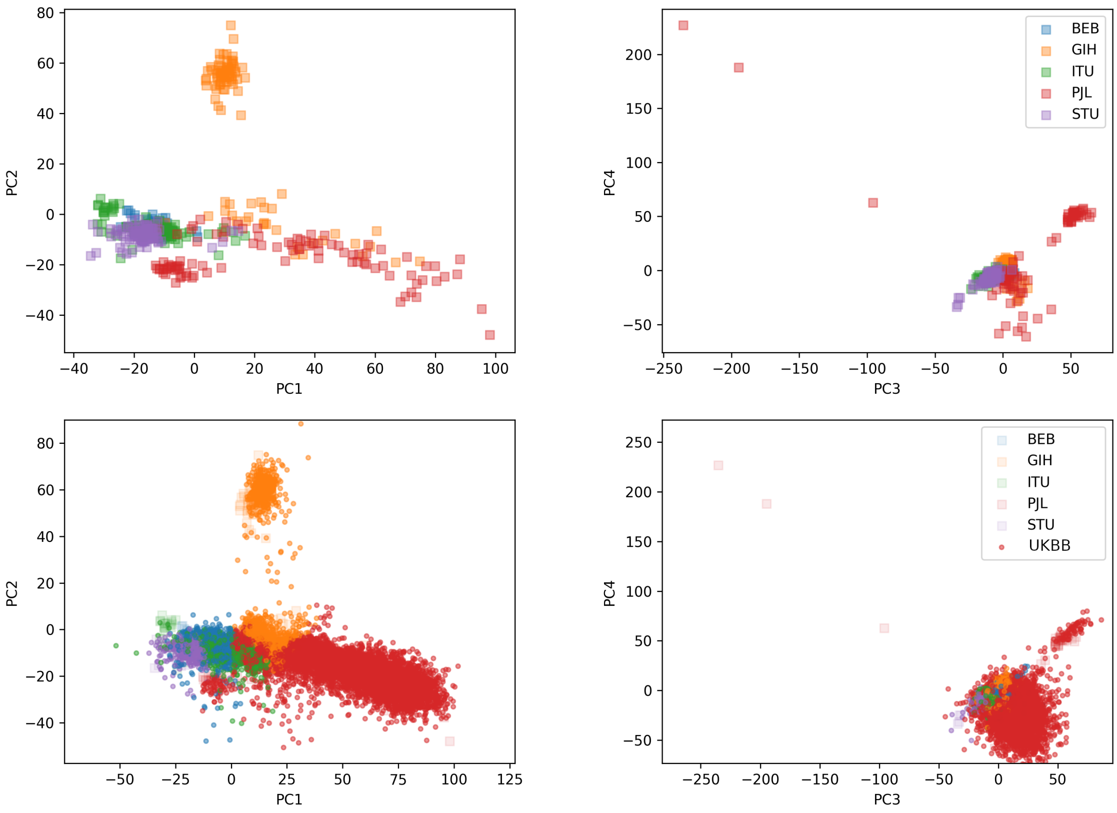


Figure S9: Top plots: Reference PC score of the 487 South Asian samples from the 1000 Genome data. Three outliers have been removed. Bottom plots: PC scores of the 10250 South Asian study samples from the UK Biobank data predicted by OADP. Different colors represents the predicted ancestry of the UK Biobank samples.

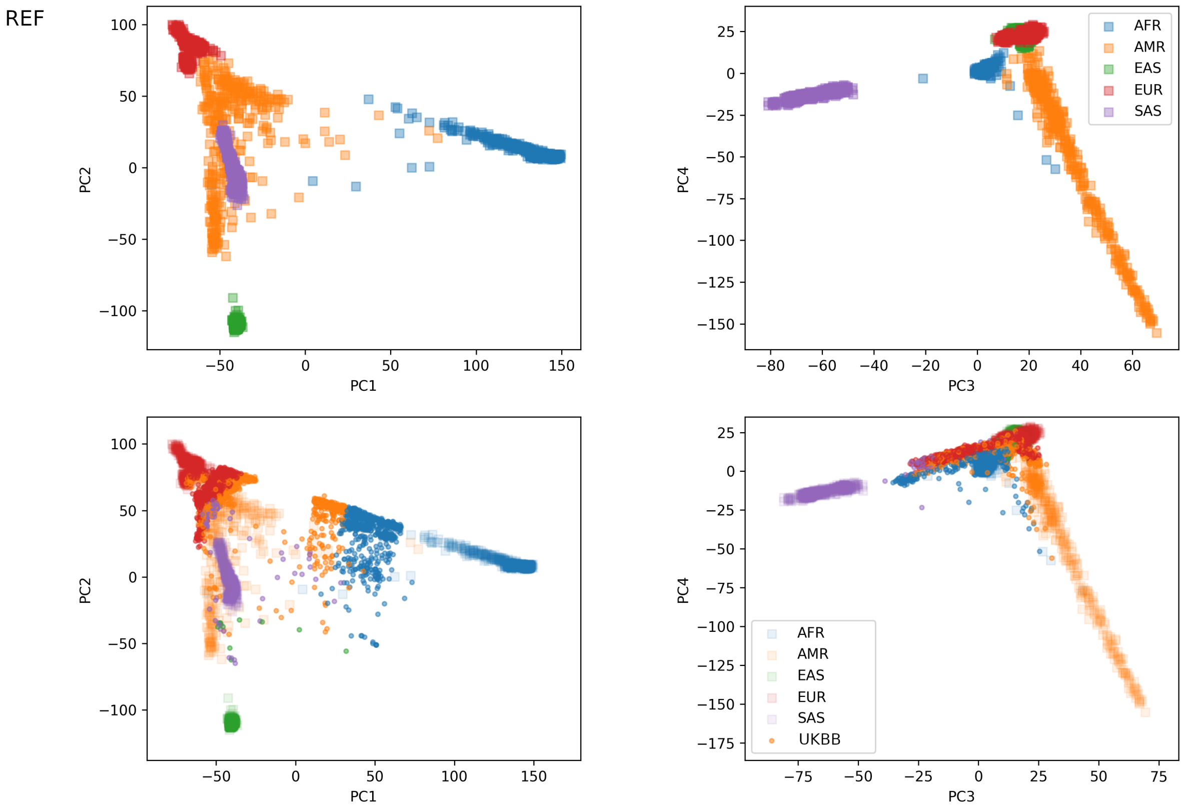


Figure S10: Top plots: Reference PC score of the 2492 global samples from the 1000 Genome data. Bottom plots: Circles are the PC scores of the 3424 study samples of mixed ancestry memberships from the UK Biobank data predicted by OADP. Mixed samples are defined to be those whose highest population prediction probability is 0.875 or less by the nearest neighbor method with weights inversetly proportional to the distances. The color of each circle represents the population with the greatest prediction probability for that sample.

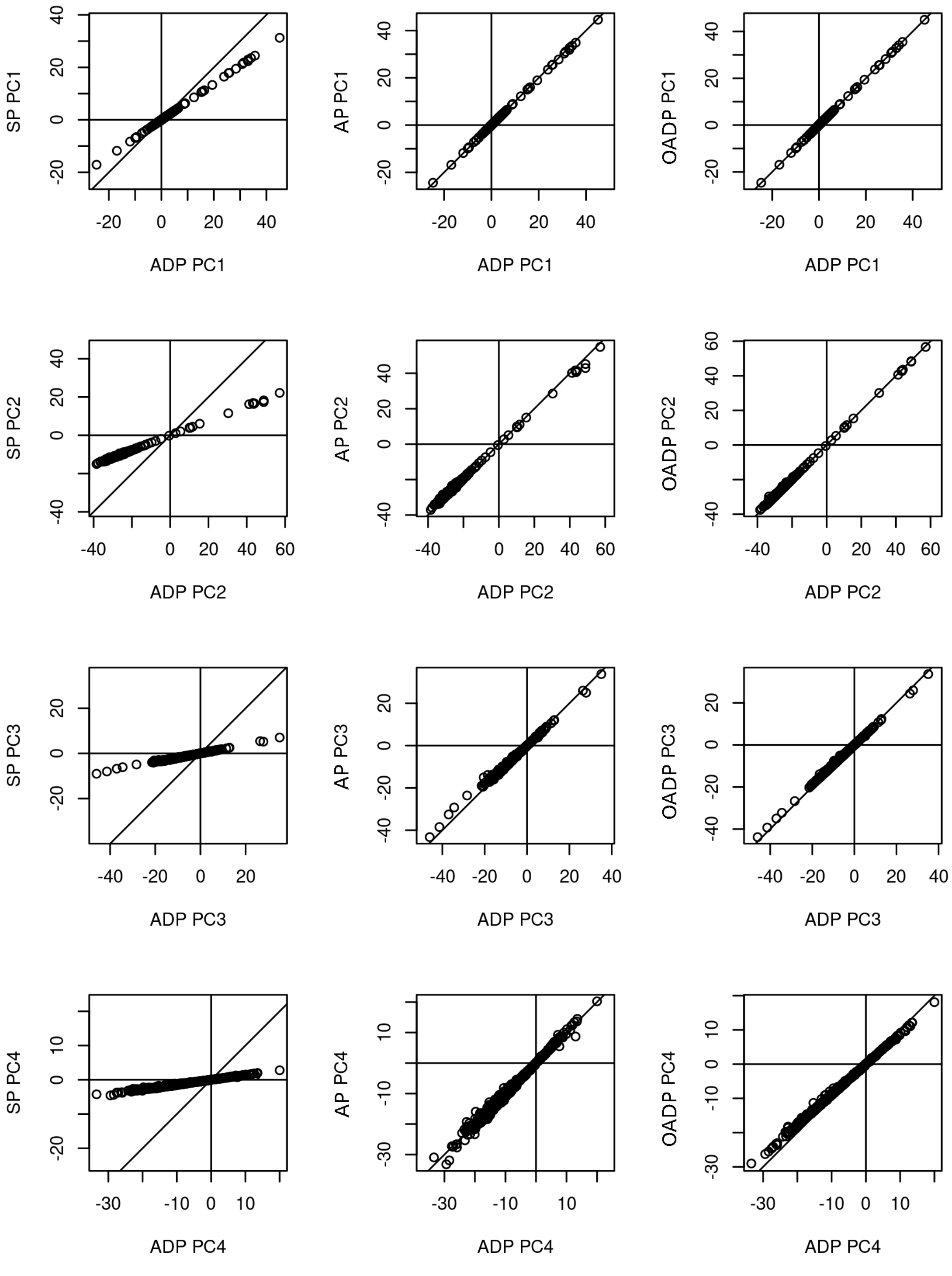


Figure S11: Comparison of the predicted PC scores of SP, AP, and OADP to ADP for the 500 randomly selected European UK Biobank study samples. The 498 European 1000 Genomes samples are used as the reference set.
